# AI-Driven and 3D-Bioprinted New Approach Methodology (NAM) Identifies NEO100 as Potent Ultrasound-Activated Therapeutic for Primary and Metastatic Brain Tumors

**DOI:** 10.1101/2025.11.25.690556

**Authors:** Rudrajit Majumder, Priyankan Datta, Sreejesh Moolayadukkam, Brooke Nakamura, Ulises Izquierdo, Radhika Joshi, Saman Sedighi, Thomas C Chen, Josh Neman

## Abstract

Primary and metastatic brain tumors are among the deadliest and treatment-resistant cancers, mainly because of their inherent resistance to chemoradiation and limited drug delivery across the blood–brain barrier (BBB). Identifying molecules that can cross the BBB and serve as sonosensitizers is crucial for developing noninvasive, targeted therapies such as sonodynamic therapy (SDT). To overcome the limitations of traditional low-throughput screening, a New Approach Methodology (NAM) was developed, starting with AI-driven molecular discovery. A positive–unlabeled neural network, trained on over 200 molecular descriptors, was used to predict compounds likely to respond to focused ultrasound (FUS), penetrate the BBB, and mediate therapeutic sonodynamic activity. While AI helps prioritize potent candidates, effective SDT also depends on clinically scalable FUS delivery. Low-intensity FUS enables noninvasive activation of small molecules in the brain and has been validated in several clinical settings. Therefore, AI-guided predictions were combined with ultrasound-based sonodynamic testing using parameters relevant to human therapy. A major challenge in SDT development is the lack of rapid, physiologically relevant tumor models, as traditional 3D organoids take weeks to mature, delaying validation. To address this, a magnetic bioprinting platform was used to produce uniform 3D tumor spheroids within hours, enabling high-throughput screening of SDT. These spheroids replicate key microenvironmental gradients, supporting reliable ultrasound-driven cytotoxicity testing. Through this combined NAM pipeline, AI identified NEO100—an ultrapure, pharmaceutical-grade perillyl alcohol currently in Phase 2a clinical trials—as a promising sonosensitizer candidate. These predictions were validated in rapidly 3D bioprinted tumor models representing glioblastoma, Group 3 pediatric medulloblastoma, meningioma, and breast-to-brain and lung-to-brain metastases. In all tumor types, ultrasound activation significantly increased NEO100’s cytotoxicity. Given its established safety profile in humans and ability to cross the BBB, NEO100 demonstrates how integrating AI-based molecular discovery, accelerated 3D bioprinting, and clinically relevant ultrasound parameters can rapidly advance precision sonodynamic therapies for various primary and metastatic brain cancers.

## Introduction

Patients with primary and metastatic brain tumors continue to face poor clinical outcomes, emphasizing the urgent need for more effective treatments. Median survival for brain metastases from breast and lung cancers is around six months [1–5], and glioblastoma still has a poor prognosis despite aggressive treatments like surgery, radiation, and temozolomide [6]. Pediatric medulloblastoma and high-grade meningiomas also show high recurrence rates and significant long-term complications [7, 8]. Progress in treating these tumor types is primarily hindered by biological and anatomical barriers that limit the delivery and effectiveness of systemic therapy treatments.

A major challenge is the difficulty in identifying small molecules that can cross the blood–brain barrier (BBB) and blood-tumor barrier (BTB), regulate relevant tumor biology [9], and be activated with spatial precision. These limitations have increased interest in sensitizer-based therapies, such as sonodynamic therapy [10, 11]. SDT uses low-intensity focused ultrasound (LIFU) to activate sensitizer molecules, producing reactive oxygen species (ROS)-mediated oxidative stress that results in targeted tumor cell death. However, the success and application of SDT rely on the molecular and electronic characteristics of sensitizers. Although knowledge of the physicochemical factors that influence photo-and sonosensitivity has improved, dependable predictive design principles remain lacking [10, 12].

Artificial intelligence (AI), offers a way to overcome this obstacle. Advanced cheminformatics and machine learning techniques can analyze complex structural, physicochemical, and electronic features linked to therapeutic response [10]. However, discovering ultrasound-activatable small molecules, or sonosensitizers, involves unique challenges: positive examples are scarce, negative examples are uncertain, and the biological factors governing acoustic activation are not fully understood. To address these issues, a positive–unlabeled (PU) neural network framework was developed to identify molecular fingerprints associated with sensitizers, utilizing over 200 RDKit descriptors and quantum-derived features [13, 14]. This method predicts which compounds are likely to respond to FUS, cross the BBB, and produce cytotoxic ROS.

While AI can prioritize candidate sonosensitizers, the success of sonodynamic therapy also depends on reliable, clinically feasible energy delivery [15]. Advances in LIFU have enabled noninvasive intracranial targeting. FUS can safely transmit acoustic energy through the skull, as demonstrated by FDA-approved indications for essential tremor and Parkinsonian tremor [16]. Moreover, low-intensity pulsed ultrasound (LIPU) can transiently open the BBB, and modern MRI-guided FUS systems enable precise sub-millimeter targeting [17–19]. Unlike photodynamic therapy, which is limited by shallow light penetration, SDT enables the activation of small-molecule sensitizers deep within the brain, positioning FUS as a transformative platform for non-invasive treatment.

Even with AI predictions and advanced FUS delivery, SDT development remains limited by biological modeling challenges. Traditional 2D cultures can’t replicate tumor hypoxia, extracellular matrix composition, or microenvironmental gradients [20–23], which are crucial for sensitizer activity. Animal models, though physiologically accurate, are too slow, expensive, and low throughput for quick screening or personalized testing. To overcome these issues, we developed a magnetic-field–guided 3D bioprinting system that produces uniform, physiologically relevant tumor spheroids within hours — much faster than conventional organoids, which can take weeks or months to produce [24–28]. These printed spheroids mimic key microenvironmental features of tumors, such as oxygen gradients and hypoxia [12], which influence treatment, allowing for high-throughput experimental validation of AI-selected sonosensitizers.

In this study, we combine PU-learning AI modeling, clinically scalable ultrasound activation, and rapid 3D bioprinting to develop a unified New Approach Methodology (NAM) for discovering and validating sonosensitizers. We apply this integrated platform to NEO100, an ultrapure, pharmaceutical-grade formulation of perillyl alcohol currently in Phase 1/2 clinical trials for primary and metastatic brain tumors [29]. AI modeling identifies NEO100 as a strong candidate sensitizer, predicting ultrasound responsiveness, BBB penetration, and ROS-generating capability. We confirm these predictions in magnetically bioprinted spheroids from glioblastoma, medulloblastoma, meningioma, and breast-to-brain and lung-to-brain metastases. Ultrasound activation significantly enhances NEO100’s cytotoxicity across all tumor types, enabling effective tumor cell killing at lower doses and shorter exposure times **(Figure 1).**

**Figure 1:**
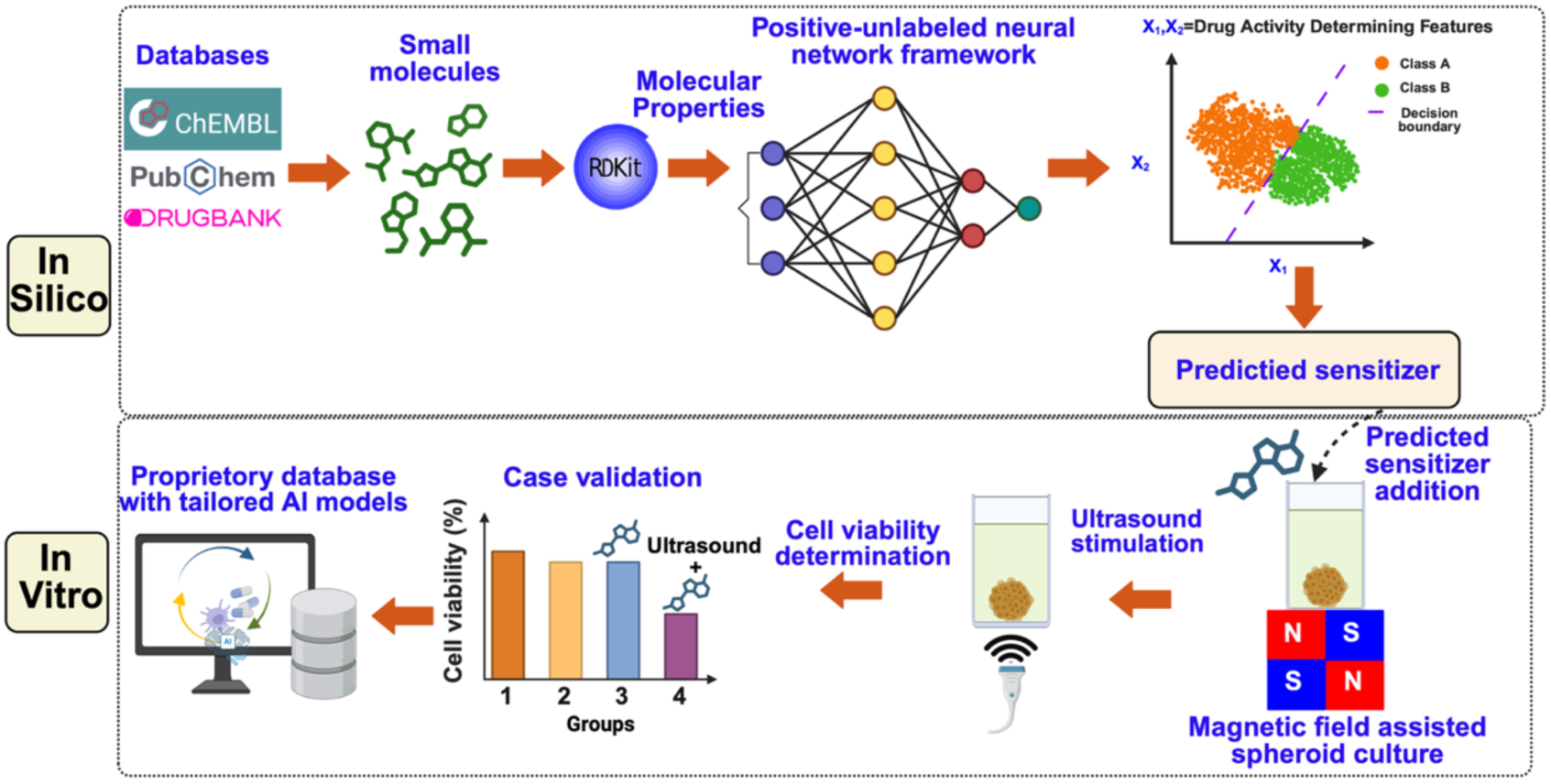
Schematic overview of the New Approach Methodology framework, consisting of two interconnected and sequential components. First, a positive–unlabeled neural network classifier conducts in silico predictions and screening of sonosensitizer candidates based on detailed molecular descriptors and quantum-derived electronic properties. Second, computationally identified candidates are validated experimentally using high-throughput, magnetic field–assisted three-dimensional cancer spheroid bioprinting platforms in vitro. This integrated translational pipeline aims to streamline and speed up the preclinical development and clinical translation of new sonodynamic therapeutic agents.

Overall, these findings demonstrate that combining AI prediction, advanced ultrasound technology, and high-quality 3D tumor models can expedite the development of clinically usable sonodynamic therapies. This complete approach presents NEO100 as a promising, noninvasive, and personalized option for patients with primary and metastatic brain tumors.

## Materials and methods

### AI-Based Positive–Unlabeled Learning Framework for Predicting Molecular Sensitizer

To enable AI-driven discovery of candidate sonosensitizers, we first assembled a curated molecular database integrating antineoplastic, immunomodulatory, and photosensitive compounds from ChEMBL[30], PubChem [31], and DrugBank [32]. Experimentally verified sensitizers were labeled as positives, whereas all other molecules were retained as unlabeled due to the lack of definitive experimental evidence demonstrating the absence of activity. Additional sensitizers from our recent systematic review of sonodynamic therapy were incorporated to broaden the positive class [10]. Using RDKit [33], we computed 212 molecular descriptors per compound. These descriptors numerically represent critical structural and physicochemical characteristics, including molecular weight, topological polar surface area, logP (a measure of lipophilicity), number of hydrogen bond donors and acceptors, atom counts, and various topological indices, which collectively inform a molecule’s potential biological behavior. This yielded a final dataset of 366 small molecules, of which 176 were confirmed sensitizers. Because unlabeled molecules inevitably contain a mixture of true negatives and previously unrecognized positives, traditional supervised learning would introduce systematic bias by forcing all unlabeled compounds into a negative class. To address this challenge, we adopted a positive–unlabeled (PU) learning framework that leverages confirmed positives alongside a large unlabeled pool to derive robust decision boundaries [13, 14]. We then trained a positive-unlabeled (PU) deep neural network (DNN) classifier implemented in TensorFlow [34], incorporating dropout and L2 regularization to prevent overfitting. The model was optimized using the non-negative PU (nnPU) loss formulation to ensure stable convergence in the absence of reliable negative samples. To assess the robustness of the class prior estimate, we performed a sensitivity analysis by varying 𝜋 from 0.1 to 1.0.

### AI Model Development

We have chosen a Deep Neural Network (DNN) as our classifier due to its theoretical capacity as a universal function approximator [35, 36]. This property guarantees that a multilayer feedforward network can approximate any function, making it highly suitable for learning complex, nonlinear relationships within data.

In a standard supervised binary classification task, the goal is to learn a mapping from an input feature matrix 𝑋 to a discrete output 𝑦, where 𝑦 ∈ {0,1}. The DNN learns this mapping by adjusting its internal parameters—weights (*w*) and biases (*b*)—to minimize a loss function that quantifies the discrepancy between the predicted probabilities 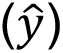 and the actual labels (𝑦).

For a conventional binary classification problem with confirmed positive and negative samples, the appropriate loss function is the Binary Cross-Entropy (BCE). The BCE loss measures the divergence between the accurate label distribution and the predicted probabilities. For a single data point, it is defined as:

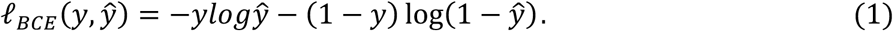

For a dataset with 𝑀 samples, this is averaged to form the total loss:

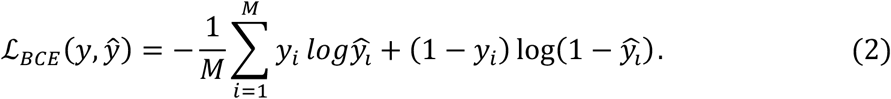

The optimization of the model parameters (*w*, *b*) is achieved via gradient descent, an iterative algorithm that minimizes the loss function [37]. The gradients of the loss with respect to each parameter are computed through the backpropagation algorithm, which applies the chain rule to propagate the error backward through the network layers [38]. This enables precise weight and bias updates, guiding the model toward a state that accurately classifies the input data.

In our specific problem, we lack reliable negative samples (*N*); our data consists only of Confirmed Sensitizers (positive samples, *P*) and Non-Confirmed molecules (unlabeled samples, *U*). The unlabeled set *U* contains both hidden positive and true negative samples. Directly applying BCE loss is inappropriate because it would incorrectly treat all unlabeled samples as negatives.

The connection between BCE and PU loss is derived from a risk minimization perspective. The expected BCE risk, 𝑅(𝑓), for a classifier 𝑓 can be expressed as:

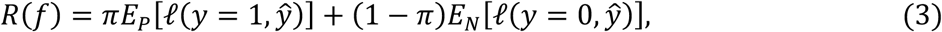

Where 𝜋 = 𝑝(𝑦 = 1) is the class prior (the proportion of true positives in the entire dataset), 𝐸_p_ and 𝐸*_N_* denote expectations over the actual positive and negative distributions, respectively. Since we have no explicit negative samples, the key insight is to represent the risk over the negative distribution as a risk over the unlabeled and positive distributions. Kiryo et al. [39] showed that the approximate relationship among these risks that yields the unbiased PU (uPU) risk estimator:

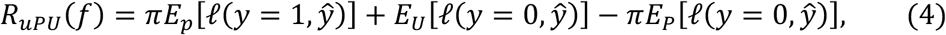

Using the BCE loss components where 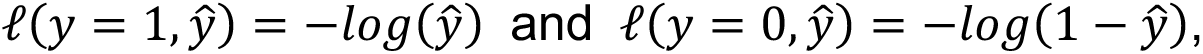 the empirical uPU loss for training is:

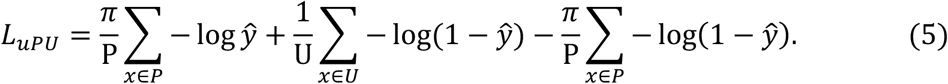

This loss is theoretically unbiased, meaning it is equivalent to training on fully labeled data. However, in practice, if the classifier becomes too confident in the unlabeled set (a common occurrence in overfitting), it leads to negative loss values and severe overfitting.

To address the instability of the unbiased PU loss, Kiryo et al. [39] proposed the non-negative PU (nnPU) loss. The modification is simple yet critical: if the empirical risk term for the unlabeled data becomes smaller than the risk term for the positives, it is clipped to zero. This prevents the loss from becoming negative and stabilizes the training process. The nnPU loss is defined as:

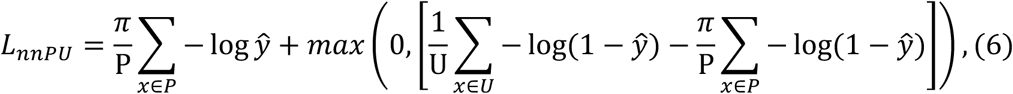

The 𝑚𝑎𝑥 operation ensures the loss component from the unlabeled data is non-negative, making the training process significantly more robust.

The PU learning framework requires an accurate estimate of the class prior 𝜋 before training. We utilized the method proposed by Elkan and Noto[40], which provides a simple and effective estimator. The core assumption of the process is that the labeled positive set *P* is selected entirely at random (SCAR) from the entire set of positive instances. The SCAR assumption introduces a key scalar constant, *c*, defined as the probability that a positive sample (𝑦 = 1) is labeled (𝑠 = 1):

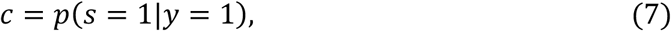

Physically, *c* represents the labeling frequency. It is the fraction of true sensitizers that have been correctly identified and included in our positive set *P*. A low *c* indicates that we have missed many sensitizers (they reside in *U*), while a high *c* suggests our positive set is relatively complete.

The Elkan & Noto method provides a simple yet elegant way to estimate *c*. The core idea is to train a probabilistic classifier, *g* (e.g., Random Forest), to distinguish the labeled positive set *P* from the unlabeled set *U*. This classifier learns to model the probability 𝑔(𝑥) = 𝑝(𝑠 = 1|𝑥), i.e., the probability that a given sample *x* is in the labeled set. The value of *c* is estimated using the trained classifier *g* and a validation set of labeled positives *P*. The estimator is the average value of *g*(*x*) for all 𝑥 ∈ 𝑃:

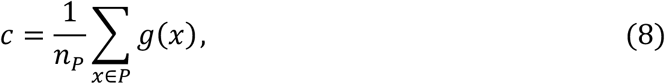

Where 𝑛_p_ is the number of labeled positive samples in the validation set. The class prior 𝜋 = 𝑝(𝑦 = 1) and the label frequency *c* are closely related to each other [40, 41]. Given a PU dataset, if one is known, the expected value of the other can be calculated as follows:

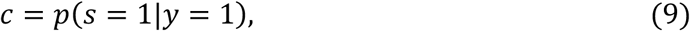

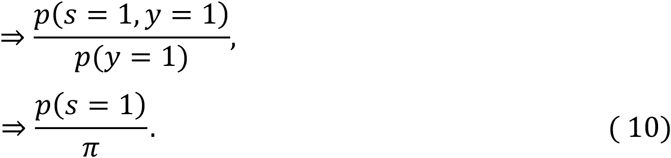

In practice, 𝑝(𝑠 = 1) is simply the empirical proportion of labeled positives in the entire dataset, 𝑛_p_⁄(𝑛_p_ + 𝑛*_U_*). Therefore, the class prior can be estimated as:

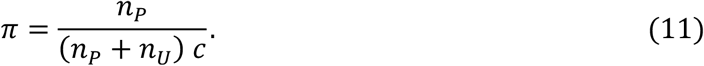

Applying this method to our dataset, we estimated the class prior 𝜋 to be 0.535. This indicates a 53.5% probability that a randomly selected molecule from the database is a true sensitizer. **Figure 2A** shows the class distributions as computed using Elkan-Noto’s method. With π determined, we proceeded to train our DNN using the robust nnPU loss framework.

**Figure 2:**
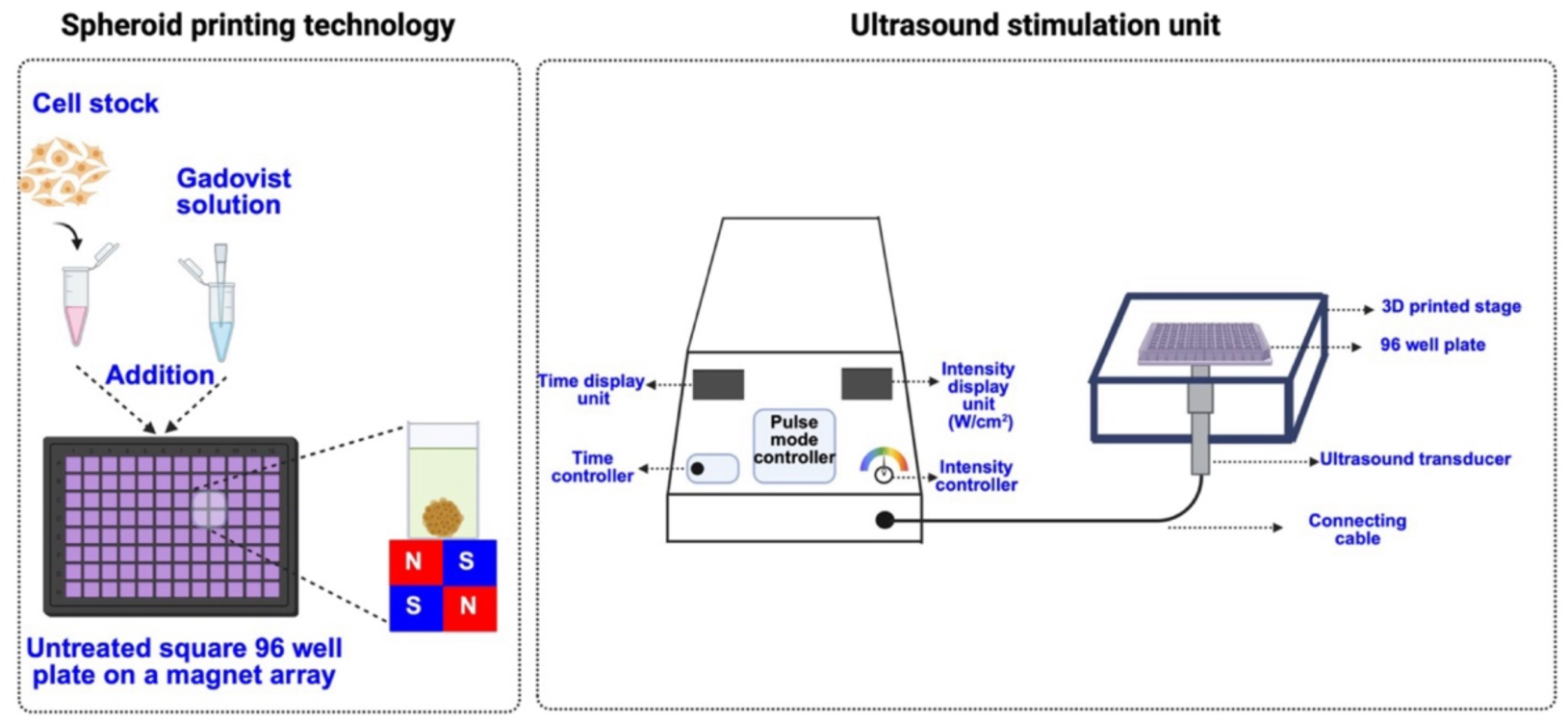
Schematic diagram of spheroid printing technology and ultrasound stimulation setup. Spheroids from five different cell lines are printed by seeding cells in culture medium containing the paramagnetic salt Gadovist®. The wells are placed on a cube magnet array and incubated at 5% CO_2_ and 37°C. After 48 hours, 96-well plates with spheroids are positioned on top of an ultrasound transducer head and stimulated from below for 5 minutes at 1 MHz, 1 W/cm², 20% duty cycle, and a 200-millisecond pulse duration.

Our Deep Neural Network (DNN) architecture was designed with one input layer, three hidden layers comprising 64, 16, and 8 neurons, respectively, and a single-node output layer. Each hidden layer was followed by a Rectified Linear Unit (ReLU) activation function to introduce non-linearity, while a sigmoid activation function was applied to the output layer to constrain predictions to a probabilistic range [0, 1]. To mitigate overfitting, dropout layers were incorporated after each hidden layer. Additionally, 𝐿_1_ kernel regularizers are employed in the hidden layers to ensure smooth gradient descent. The model was compiled using the Adam optimizer with a learning rate of 0.001. The dataset was partitioned into training (70%), validation (10%), and test (10%) sets, and stratified to preserve the class distributions in all splits.

To assess the sensitivity of our model to the class prior 𝜋 and validate the value estimated by the Elkan & Noto method, we performed a hyperparameter analysis by varying 𝜋 from 0.1 to 1.0. For each value, the model was trained five times, and the resulting recall on the test set was recorded.

### Reagents and chemicals

GLPBIO Cell Counting Kit-8 (CCK-8), Promega CellTiter-Glo (R) 3D cell viability assay (G9681), GibcoTM DPBS (14040133), GibcoTM DMEM (11995040), Trypan Blue (0.4%, Thermofisher Scientific, USA), Invitrogen ready probes cell viability imaging kit, Blue/Green (R37609), are purchased from Fisher Scientific, USA, and Dimethyl Sulfoxide (for molecular biology, D8418), Millipore Sigma, USA.Trypsin-EDTA solution (1X, 30-2101) is purchased from ATCC, USA. NEO 100TM is provided by NeOnc Technologies Holdings Inc. (NTHI) through the University of Southern California Brain Tumor Center.

### In vitro 2D cell culture

U-87 MG cell lines are purchased from ATCC, USA. Grade 3 medulloblastoma cell line D425, patient-derived grade 2 meningioma cell MN5, patient-derived breast to brain metastasis (BBM3.1), and lung to brain metastasis (LuBM5) are obtained from the University of Southern California Brain Tumor Center. U-87 MG, D425, LuBM5, and BBM3.1 cells are cultured in a complete medium of Dulbecco’s Modified Eagle Medium, supplemented with 10% Fetal Bovine Serum (Thermofisher Scientific, USA, Cat. No. A5256801), 1% Penicillin-Streptomycin (Fisher Scientific, USA), and maintained at 5% CO_2_ and 37^0^C *in vitro*. MN5 cells are cultured in a mixture (1:1 v/v) of complete advanced DMEM F12 (Thermofisher Scientific, USA, Cat. No. 12634010) and neurobasal medium (Thermofisher Scientific, USA, Cat No.21103049) supplemented with 10% FBS, 2% B-27, 1% Glutamax, and 1% Penicillin-Streptomycin (Fisher Scientific, USA), and maintained at 5% CO_2_ and 37^0^C *in vitro*.

### Magnetic Field–Guided 3D Bioprinting of Brain Tumor Spheroids and Sonodynamic Treatment

Primary (U-87 MG, D425, MN5) and metastatic brain (BBM3.1, LuBM5) tumor spheroids were generated using proprietary magnetic-field–guided 3D cell-printing technology (NeOnc Technology Holdings Inc Patent US 11,788,957), following previously published protocols [12]. Cells were harvested, counted, and dispensed into an uncoated, flat, square 96-well plate (Ibidi, Cat. No. 89621) positioned on a cube magnet array (4.5 mm × 4.5 mm × 4.5 mm). Each well received 10,000 cells suspended in culture medium supplemented with a paramagnetic salt solution (Gadovist®, Bayer). The magnetic susceptibility difference between the diamagnetic cells and the surrounding paramagnetic medium produced a net magnetic force that drove cells toward the region of lowest magnetic field strength, promoting rapid aggregation. This process yielded layer-by-layer 3D cellular constructs within hours, with timing dependent on tumor type (**Figure 3).** As previously reported, U-87 MG spheroids generated by this approach form a hypoxic core within 12 h [12], validating the physiological relevance of the printed microstructures. For sonodynamic therapy studies, 3D spheroids derived from glioblastoma, medulloblastoma, meningioma, and metastatic brain cancer cell lines were incubated for 48 hours following printing (**Figure 3**). On day 3, NEO100 was added at its IC_50_ and IC_25_ concentrations and incubated for an additional 4h. Ultrasound stimulation was delivered using a transducer (Delta Digi Sound) mounted on a custom 3D-printed stage fabricated from polylactic acid using a Prusa MK4 printer. The transducer head (3 cm diameter) was positioned beneath the well plate, and ultrasonic transmission gel (Aquasonic® 100, Parker Laboratories) was applied to minimize acoustic impedance mismatch. Ultrasound exposure was performed at 1 MHz, 1 W/cm², 20% duty factor, 200-ms pulse duration, and a total sonication time of 5 min. Spheroid viability was assessed 24 h after stimulation using ATP quantification with the CellTiter-Glo® 3D Cell Viability Assay (Promega).

**Figure 3:**
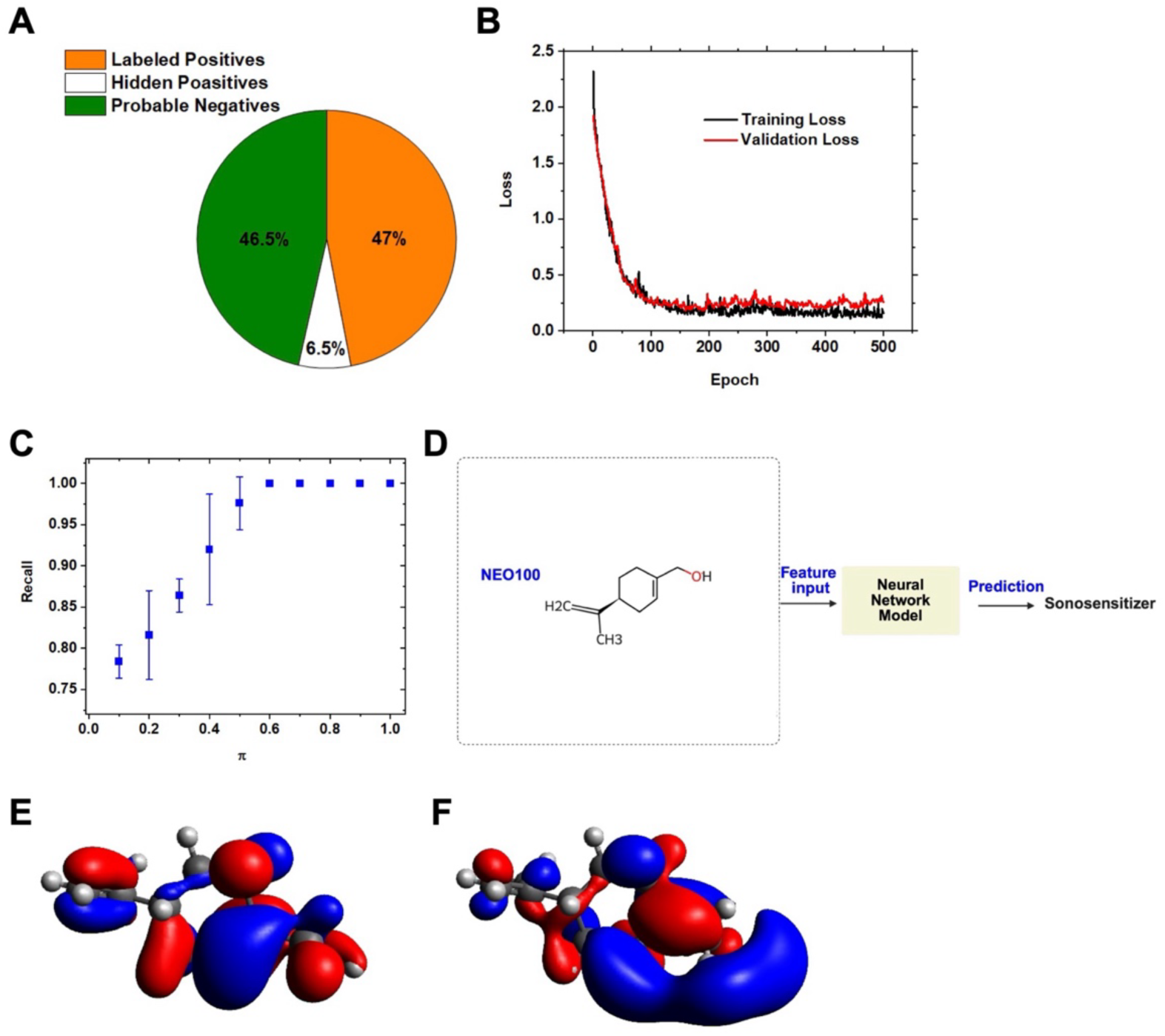
Neural Network Development, Validation, and Application for Sonosensitizer Prediction. (A) Pie chart showing the dataset composition based on the Elkan & Noto class prior (𝜋 = 0.535). The unlabeled set is divided into hidden positives and likely negatives, indicating a PU learning scenario. (B) Training and validation loss curves of the DNN model over 500 epochs, showing convergence with minimal overfitting. (C) Sensitivity analysis of the class prior (𝜋). Model recall is measured across various π values (mean ± standard deviation, n = 5 runs). (D) Workflow diagram of the trained DNN model used to predict a candidate molecule, NEO100, as a sonosensitizer. (E) HOMO at-8.690 eV and (F) LUMO at 2.162 eV for NEO100. The energy gap between LUMO and HOMO for NEO100 is 10.852 eV. The positive and negative phases of orbital wavefunctions are represented in red and blue.

### 3D-bioprinted spheroid area calculation

Brightfield images for each cell line (n = 10) are obtained at regular intervals over 6 days using a Leica microscope to evaluate the spheroid area. The images are then processed using ImageJ. Each image is first converted into grayscale, and the Otsu threshold is applied to separate the spheroid from the background. Further morphological operations (internal gaps correction via hole filling, and small background particle exclusion using size-based filtering) are performed to refine the binary mask. The largest connected region is identified as the spheroid, and its pixel area is measured. Thereon, the spheroid area is evaluated using the Leica microscope calibration factor.

### IC_50_ determination for NEO100

IC₅₀ values for 3D tumor cultures were determined by quantifying ATP release using the CellTiter-Glo® 3D Cell Viability Assay (Promega), following the manufacturer’s protocol and procedures consistent with our earlier work [12]. Primary and metastatic brain tumor spheroids were printed at 10,000 cells per well and incubated for 48 h. On day 3, NEO100 was added at varying concentrations and incubated for 72 h. After treatment, the culture medium was replaced with fresh medium, and the CellTiter-Glo® 3D reagent was added at a 1:1 (v/v) ratio. Plates were incubated in the dark for 25 min to ensure complete cell lysis and stabilization of the luminescent signal. The resulting supernatant from each well was transferred to an opaque, white, tissue culture–treated 96-well plate (353296, Becton Dickinson Labware, USA) to minimize signal attenuation. Luminescence was recorded using a Synergy H4 plate reader (BioTek) with an integration time of 0.01 s. Cell viability and IC₅₀ calculations were performed using GraphPad Prism. All compounds, including NEO100, were administered as single doses in each experiment.

### Confocal imaging

Confocal images are taken 48 hrs. after cell seeding. For this purpose, printed spheroids are stained with NucBlue® Live and NucGreen® Dead reagents according to the manufacturer’s protocol, and images are captured using a Leica microscope. Briefly, two drops of NucBlue® Live and NucGreen® Dead reagents are added to the spheroid culture medium, and the mixture is incubated at 5% CO_2_ and 37 0C for 30 minutes. The live reagent stains only the live cell nuclei and is detected with a standard DAPI filter (excitation/emission maxima: 360/460 nm). The dead reagent stains only the nuclei of dead cells with compromised plasma membranes and is detected with a standard FITC (green) filter set (excitation/emission maxima: 504/523 nm). The captured images are further processed using ImarisViewer 10.2.0.

### RNA Isolation and qPCR Analysis

Cells were harvested either by scraping when suitable or by trypsinization for 3-5 minutes at 37°C, then neutralized in media containing 10% FBS, and centrifuged at 360 rcf for 5 minutes. The resulting cell pellet was either processed immediately for RNA extraction or frozen for later use in qPCR (performed in triplicate per sample), as previously described [2]. All primers used for qPCR analysis were purchased from IDT.

### Statistical analysis

Data are expressed as mean ± standard deviation. The one-way ANOVA tests for significant differences among groups; P < 0.05 is considered statistically significant. All experiments are performed at least in triplicate.

## Results

### AI-Based PU Modeling Identifies NEO100 as a High-Confidence Sonosenitizer with Favorable Quantum-Mechanical Properties

We applied a positive–unlabeled (PU) learning strategy to address the lack of experimentally verified negative controls in sensitizer discovery. Estimation of the proportion of true sensitizers in the full molecular library—the class prior 𝜋 —was performed using the Elkan–Noto method [40]. This analysis yielded a class prior of 𝜋 = 0.535 (**Figure 3A**), indicating that more than half of the curated molecules are likely to exhibit sensitizing activity. This finding reflects the dataset’s chemical diversity and provides a critical parameter for PU-based model optimization.

The deep neural network (DNN) classifier was trained using the non-negative PU (nnPU) loss function to ensure stable learning in the absence of confirmed negative samples. The model, implemented in TensorFlow [34], was trained for 500 epochs with smooth convergence and clear separation between training and validation loss (**Figure 3B**). When evaluated on an independent test set, the classifier achieved a recall of 97.6% for confirmed sensitizers, demonstrating excellent sensitivity and reliable recovery of known positives.

To evaluate the robustness of the class prior estimate, we conducted a sensitivity analysis over a wide range of priors (𝜋 = 0.1–1.0). When π exceeded approximately 0.60, the model collapsed into an all-positive prediction state, with output probabilities reaching saturation at 1.0 for all inputs—a known failure mode of nnPU classifiers when 𝜋 overestimates the actual class distribution (**Figure 3C**). Conversely, the empirically derived 𝜋 = 0.535 avoided this problematic regime while maintaining high recall. This operating point thus offers a biologically and statistically justified prior that allows the classifier to correctly identify true sensitizers while still discriminating against negatives.

Using the optimized framework, we next evaluated the predicted sonosensitizing potential of NEO100, an ultrapure pharmaceutical-grade formulation of perillyl alcohol currently undergoing clinical testing. The trained DNN classified NEO100 as a sensitizer with a confidence of 99.8% ± 0.008 (**Figure 3D**), ranking it among the top predicted candidates in the curated library and providing strong computational evidence to support its experimental assessment as a focused ultrasound–responsive therapeutic.

To complement the machine-learning predictions, we also assessed the quantum-mechanical properties of NEO100 using time-dependent density functional theory (TD-DFT) calculations. We employed the ωB97X functional with the def2-TZVP basis set on the ORCA computational platform. The highest occupied molecular orbital (HOMO) and lowest unoccupied molecular orbital (LUMO) energies for NEO100 were found to be −8.690 eV and 2.162 eV, respectively (**Figure 3E, F**), resulting in an energy gap of 10.852 eV. The positive and negative phases of the orbital wavefunctions are indicated in red and blue, respectively. Additionally, the excited-state energies, spin–orbit coupling (SOC), and intersystem crossing rates (K_ISC) show that NEO100 exhibits favorable excited-state properties consistent with its predicted sensitizer activity (**Table 1**).

**Table 1.** A summary of the TD-DFT-based electronic properties calculations for NEO100.

| **Molecule** | **Lowest singlet**<br>**state energy**<br>**(eV)** | **Lowest triplet**<br>**state energy**<br>**(eV)** | **Maximum**<br>**SOC (cm<sup>-1</sup>)** | **Dominant**<br>**$K_{ISC}$ (s<sup>-1</sup>)** |
| --- | --- | --- | --- | --- |
| NEO100 | 6.377 | 2.741 | 10.47 | $1.57 \times 10^{10}$ |

Together, these results emphasize the two main challenges addressed by our computational framework: (i) the feasibility of creating large-scale molecular descriptor datasets using RDKit while avoiding the high computational cost of full DFT-derived descriptor libraries, and (ii) the difficulty of relying on traditional supervised learning due to the limited availability of experimentally confirmed negative sensitizers. By utilizing PU learning—which can operate solely on confirmed positive and unlabeled examples—the model effectively identifies both known and previously unknown sensitizers, including NEO100. This combined approach provides a scalable basis for AI-driven discovery of candidate sonosensitizers for subsequent ultrasound testing.

### Morphological and Growth Characterization of 3D-Bioprinted Primary and Metastatic Brain Tumor Spheroids

Accurate evaluation of candidate sonosensitizers requires physiologically relevant 3D tumor models that replicate core features of the tumor microenvironment, including proliferative zoning, compaction, and necrotic core formation. To ensure that our magnetically bioprinted spheroids provide biologically meaningful platforms for downstream sonodynamic testing, we first examined their morphology, growth kinetics, and viability across primary and metastatic brain tumor cell types.

Phase-contrast imaging at 48h post-printing revealed distinct morphological signatures associated with each tumor subtype (**Figure 4A**). D425 medulloblastoma spheroids formed spheres with a dense, optically thick core and a prominent proliferative outer rim. In contrast, U-87 MG glioblastoma spheroids exhibited smooth contours and a looser structure, with single cells and small aggregates dispersed around the spheroid perimeter. LuBM5 lung-to-brain metastasis spheroids displayed a similar architecture. BBM3.1 breast-to-brain metastasis spheroids showed the highest central optical density among all models, suggesting early necrotic core formation, accompanied by a thinner proliferative rim relative to D425 and LuBM5 spheroids. MN5 spheroids exhibit a compact structure with a dense core, along with a thin proliferative rim as similar to the BBM3.1 spheroids.

**Figure 4:**
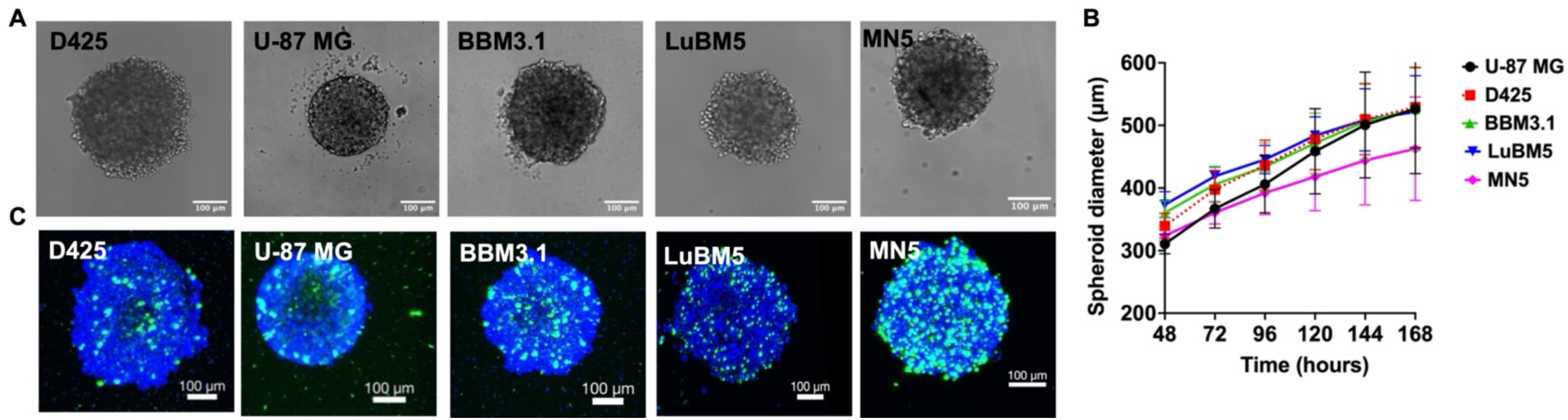
Morphology, growth kinetics, and viability of magnetically 3D-bioprinted primary and metastatic brain tumor spheroids. (A) Phase-contrast images of 3D spheroids generated using magnetic field–guided bioprinting, shown 48 hours after printing for medulloblastoma (D425), glioblastoma (U-87 MG), breast-to-brain metastasis (BBM3.1), lung-to-brain metastasis (LuBM5), and meningioma (MN5). (B) Changes in spheroid diameter over time for U-87 MG, D425, BBM3.1, and LuBM5 spheroids. (C) Confocal fluorescence images of D425, U-87 MG, BBM3.1, and LuBM5 spheroids at 48 hours post-printing. Spheroids were stained with NucBlue® Live and NucGreen® Dead ReadyProbes®, then imaged using DAPI and FITC channels on a Leica confocal microscope. All samples used identical z-stack spacing (4.28 μm). Z-stack images were reconstructed and processed using ImarisViewer software.

Spheroid expansion was quantified over six days using the Gompertz growth model [42], which captured the asymmetric sigmoidal behavior characteristic of diffusion-limited 3D tumor growth (**Figure 4B**). D425 spheroids exhibited the highest growth rate constant (0.012 h⁻¹) and the shortest doubling time (55 h), consistent with their robust proliferative rim and rapid diameter enlargement. LuBM5 spheroids displayed the next-highest growth rate (0.011 h⁻¹), accompanied by early compaction due to strong cell–cell adhesion, resulting in a faster transition to the plateau phase. U-87 MG spheroids exhibited weaker aggregation and slower expansion (0.009 h⁻¹). MN5 spheroids show a growth rate constant of 0.009 h^-1^, while BBM3.1 spheroids displayed the lowest growth rate constant (0.007 h⁻¹), in line with their dense optical cores and reduced peripheral proliferation. Confocal fluorescence imaging showed viable tumor cells with small, localized necrotic centers across all models (**Figure 4C**).

To assess whether magnetic 3D bioprinting alters tumor-related transcriptional programs, we examined the expression of key tumor suppressors, oncogenes, and epigenetic regulators—including *TP53*, *PTEN, KRAS, MTOR, HDAC3*, and *DNMT3B*—in U-87 MG glioblastoma, D425 medulloblastoma, and BBM3.1 breast-to-brain metastasis (Figure 5A–C). The 3D-printed spheroids maintained levels of several oncogenic and metabolic regulators (*KRAS, MTOR*) comparable to control conditions. Tumor suppressors *TP53* and *PTEN* were also preserved in the 3D-printed spheroids, indicating reduced stress-induced suppression compared to naturally aggregated spheroids. Similarly, epigenetic regulators *HDAC3* and *DNMT3B* showed consistent regulation, with expression maintained in 3D-printed spheroids. Overall, these results demonstrate that 3D-printed spheroids sustain physiologically relevant expression of tumor suppressors, oncogenic drivers, and epigenetic regulators compared with naturally formed spheroids, while also capturing the architectural benefits of 3D culture. This highlights magnetic bioprinting as a robust platform for modeling tumor biology and testing therapeutic responses in a controlled, reproducible 3D microenvironment.

**Figure 5:**
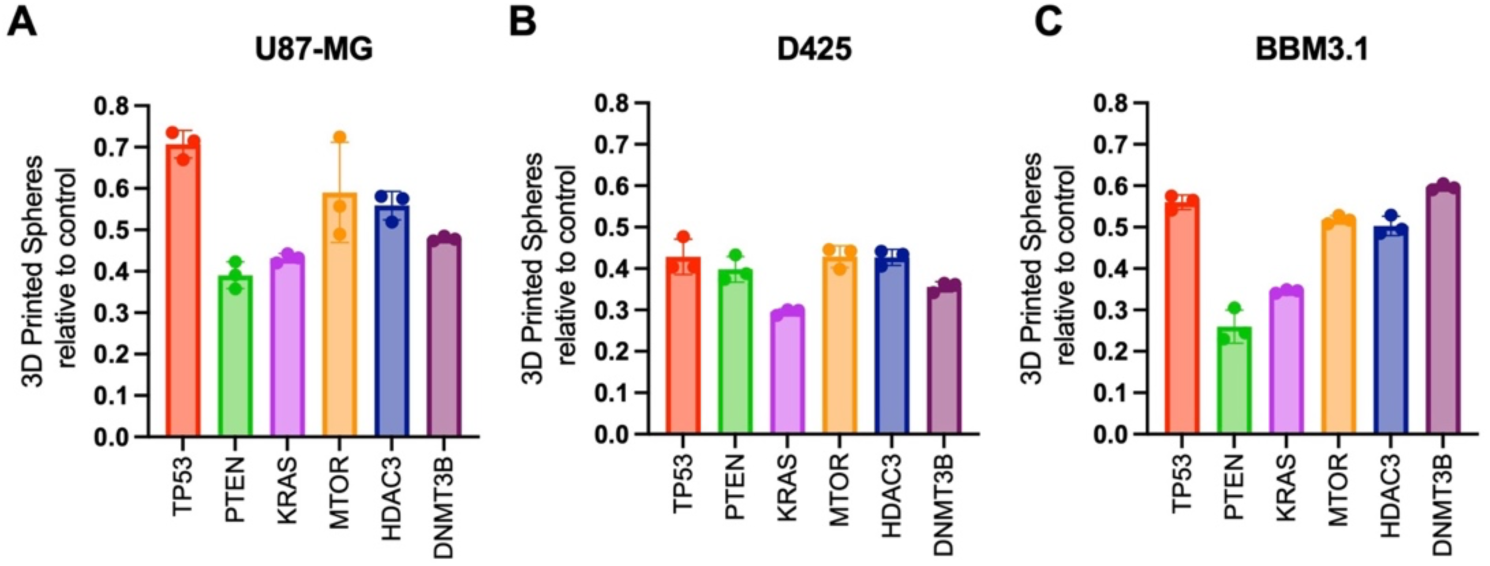
**Expression of oncogenic, tumor-suppressive, and epigenetic regulators in 3D-bioprinted spheroids compared to 2D controls. (**A–C) Quantitative PCR analysis showing expression levels of key tumor suppressors (*TP53, PTEN*), oncogenic drivers (*KRAS, MTOR*), and epigenetic regulators (*HDAC3, DNMT3B*) in 3D-bioprinted tumor spheroids relative to matched controls. Bars indicate mean ± SD (n = 3).

Together, these results validate that the magnetically bioprinted spheroids recapitulate key morphological and kinetic hallmarks of in vivo tumors—including compaction behavior, proliferative zoning, and necrosis—supporting their use as robust, physiologically relevant platforms for evaluating sonodynamic responses to NEO100.

### Ultrasound significantly enhances the cytotoxic activity of NEO100 across primary and metastatic brain tumor spheroids

To determine whether ultrasound augments the antitumor efficacy of NEO100, we quantified spheroid viability across five 3D-bioprinted brain tumor models following treatment with NEO100 alone or in combination with ultrasound stimulation.

We first quantified the cytotoxic potency of NEO100 across 3D-bioprinted primary and metastatic brain tumor spheroids (**Figure 6**). These IC₅₀ measurements revealed tumor type–specific differences in baseline sensitivity to NEO100 and established the dosing framework for subsequent sonodynamic therapy experiments. In all subsequent studies, the IC₅₀ and IC₂₅ concentrations derived from these assays were used to evaluate ultrasound-enhanced cytotoxicity. Next, US responses were evaluated across five 3D-bioprinted brain tumor models using experimentally derived IC₂₅ and IC₅₀ concentrations of NEO100 (**Figure 7A–E**).

**Figure 6:**
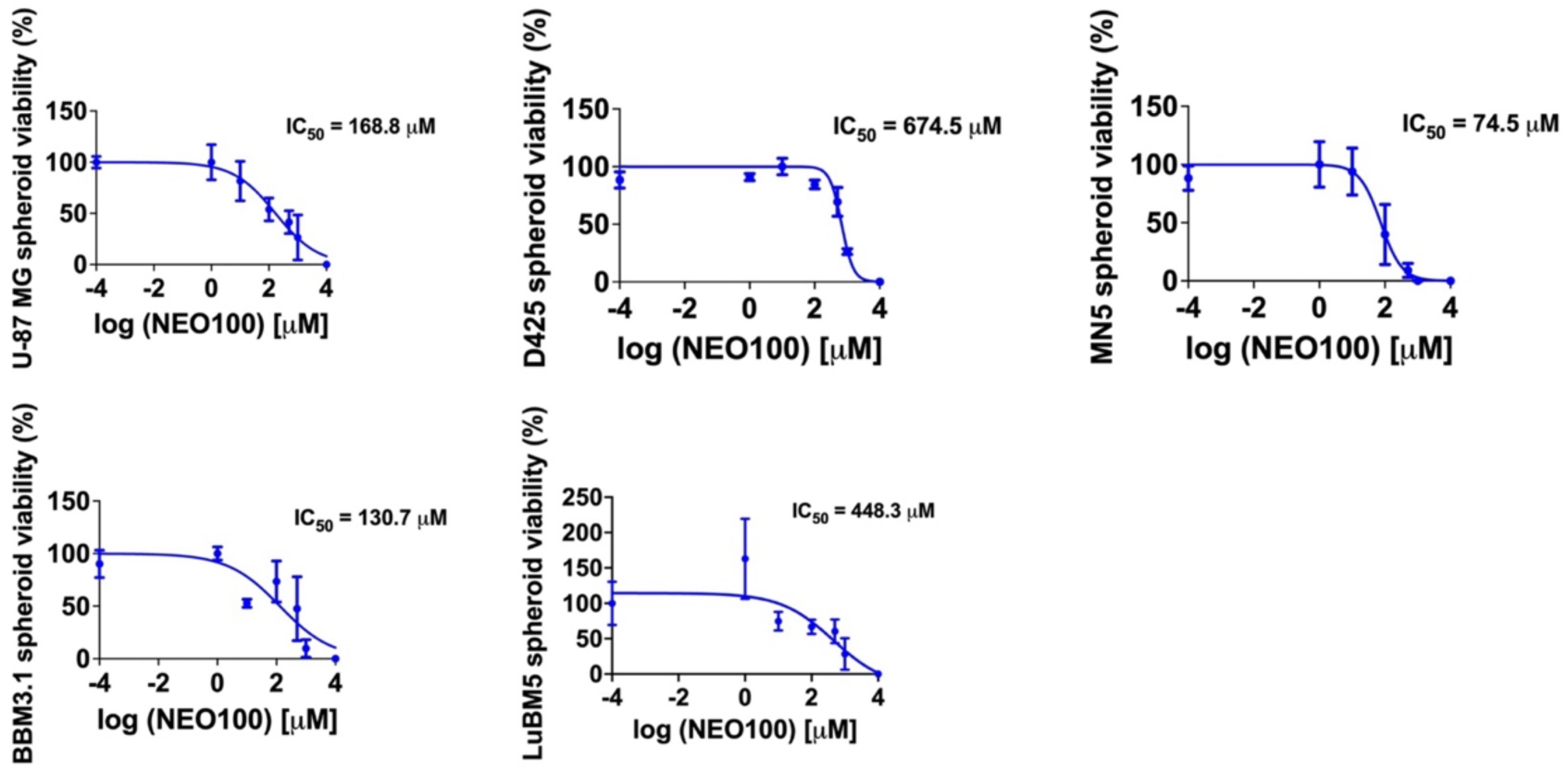
I**C₅₀ values of NEO100 across 3D-bioprinted primary and metastatic brain tumor spheroids.** IC₅₀ values of NEO100 measured in 3D-bioprinted spheroids derived from primary brain tumors (D425 medulloblastoma, U-87 MG glioblastoma, MN5 meningioma) and metastatic brain tumors (BBM3.1 breast-to-brain metastasis, LuBM5 lung-to-brain metastasis).

**Figure 7:**
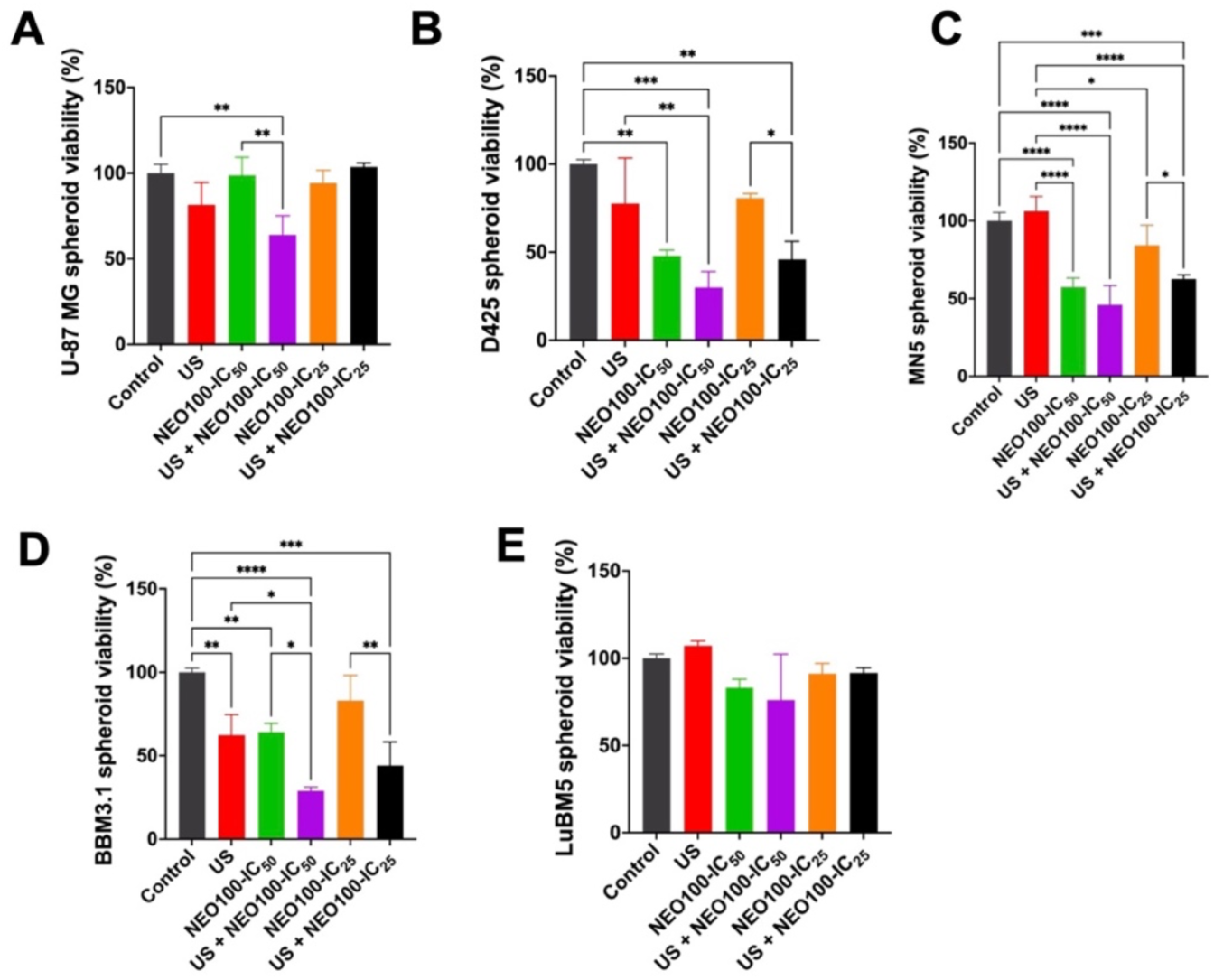
Sonodynamic therapy with NEO100 increases cytotoxicity in primary and metastatic brain tumor spheroids. (A–E) Cytotoxicity analysis of 3D-bioprinted spheroids derived from glioblastoma (U-87 MG), medulloblastoma (D425), meningioma (MN5), breast-to-brain metastasis (BBM3.1), and lung-to-brain metastasis (LuBM5) following treatment with NEO100 at IC₂₅ or IC₅₀ doses, focused ultrasound alone (US), or combined NEO100 + US. Ultrasound significantly enhances NEO100 cytotoxicity across all models in a dose-dependent manner. Statistical analysis was conducted using one-way ANOVA with Tukey’s multiple comparisons test (n = 3). *p<0.05, **p<0.0021, ***p<0.0002 ****p<0.0001.

In U-87 MG glioblastoma spheroids (**Figure 7A**), NEO100 at IC₂₅ and IC₅₀ produced moderate reductions in viability compared to control; however, combining ultrasound with NEO100 resulted in significantly enhanced cytotoxicity. Moreover, tumor viability decreased significantly under NEO100 IC₅₀ combined with US than IC₂₅ combined with US, confirming dose-dependent potentiation of NEO100 by ultrasound.

D425 medulloblastoma spheroids also demonstrated a strong overall response (**Figure 7B**). Specifically, NEO100 IC₂₅ alone modestly reduced viability, whereas NEO100 IC₂₅ combined with US produced a significant decrease in viability. NEO100 IC₅₀ with the addition of ultrasound produced an even significantly greater cytotoxic effect, showing that medulloblastoma is highly responsive to ultrasound-mediated activation of NEO100.

In MN5 meningioma spheroids (**Figure 7C**), NEO100 IC₂₅ had limited cytotoxicity, but NEO100 IC₂₅ combined with US produced a significant reduction in tumor cell viability. This trend was enhanced at NEO100 at IC₅₀, where ultrasound led to a highly significant decrease, indicating strong ultrasound sensitization even in this slower-growing primary tumor type.

BBM3.1 breast-to-brain metastatic spheroids exhibited notable dose-dependent responses (**Figure 7D**). NEO100 IC₂₅, combined with US, significantly reduced tumor viability. Furthermore, NEO100 IC₅₀ combined with US displayed a significantly greater reduction, demonstrating substantial sensitization of this metastatic model to NEO100 cytotoxicity enhanced by ultrasound. In contrast, LuBM5 lung-to-brain metastatic spheroids (**Figure 7E**) showed no appreciable cytotoxicity to NEO100, with or without ultrasound, relative to untreated controls.

Collectively, these findings demonstrate that NEO100 exhibits robust ultrasound-responsive cytotoxicity in both primary and metastatic brain tumor spheroids, confirming a strong, reproducible sonodynamic effect. These results highlight NEO100’s promise as a broadly effective sonodynamic therapeutic agent and indicate that intrinsic tumor-specific biological properties can modulate the degree of responsiveness to ultrasound-enhanced treatment.

## Discussion

The present study introduces an integrated New Approach Methodology that combines AI-driven compound discovery, rapid magnetic 3D bioprinting, and clinically translatable ultrasound to expedite the identification and validation of sonodynamic therapies for primary and metastatic brain tumors. This development aligns directly with the ongoing National Institutes of Health (NIH)-wide initiative to reduce or replace in vivo animal testing by adopting physiologically relevant human 3D tissue models, high-content screening platforms, and computational prediction tools. As NIH emphasizes advanced organoid technologies and in vitro models for therapeutic screening, our NAM presents a next-generation alternative that generates uniform, biologically mature spheroids within days rather than weeks, thereby overcoming the primary time constraints associated with traditional organoids.

Our NAM directly addresses several NIH priorities: A) Reducing animal testing—by producing physiologically relevant spheroids in hours rather than weeks, the platform substitutes for lengthy xenograft procedures and decreases animal use without compromising biological accuracy. B) Promoting organoid adoption—rapid 3D bioprinting shortens organoid development from weeks to hours, enabling real-time screening, iterative refinement, and patient-specific therapeutic testing. C) Supporting computational–experimental integration—merging AI-driven molecular discovery with human 3D tumor models aligns with the NIH’s NAMs initiative to enhance scientific rigor and predictive accuracy for translational research. D) Accelerating oncology drug discovery—high-throughput SDT screening now becomes feasible, permitting swift evaluation of sensitizers, acoustic parameters, and drug combinations within clinically relevant tumor microenvironments.

Our AI results show that combining positive–unlabeled (PU) learning with a custom descriptor library offers a significantly better strategy for identifying therapeutics capable of functioning in the brain’s uniquely restrictive environment. Traditional cheminformatics models struggle to classify sonosensitizers because negative training examples are essentially unknown failure to demonstrate sensitization experimentally does not confirm true inactivity. By using PU learning, our model avoids the systematic bias found in typical binary classifiers and instead infers molecular patterns associated with sensitizers directly from confirmed positives and a large pool of unlabeled data. This method is especially useful for brain tumor drug discovery, where validated sensitizers are rare, and chemical space is enormous.

The strength of this framework lies in the design of the new database itself. Unlike typical drug-likeness datasets, our curated library combines over 200 RDKit descriptors with quantum-mechanical parameters—properties highly relevant to BBB penetration, ROS generation, and ultrasound responsiveness. Including HOMO–LUMO energy gap, excited-state transitions, and intersystem crossing rates ensures that the model detects electronic features directly connected to cytotoxic ROS formation, which is essential for effective sonodynamic therapy. These quantum descriptors, therefore, capture mechanistic features that traditional physicochemical metrics cannot, making the database uniquely suited for identifying agents active in deep-seated intracranial tumors.

The successful prediction of NEO100 as a high-confidence sonosensitizer highlights the effectiveness of this integrated approach. With a 99.8% predicted chance of sensitization and quantum parameters indicating strong excited-state reactivity, NEO100 ranked among the most promising candidates across the entire molecular library. Importantly, this prediction was confirmed experimentally in several primary and metastatic brain tumor models, validating both the algorithm and the database’s design principles. The close agreement between computational predictions and biological results shows that a model combining PU learning with quantum-enhanced descriptors can reliably identify therapeutics with the necessary electronic and physicochemical properties for BBB permeability, acoustic activation, and ROS-mediated cytotoxicity.

Beyond identifying NEO100, the broader implication is clear: our new AI database and PU-learning pipeline create a mechanistically informed, scalable foundation for discovering brain-penetrant, ultrasound-responsive therapeutics—something that cannot be achieved with existing drug-likeness datasets. This approach could transform early-stage CNS oncology drug discovery by enabling computational triage of chemical space, reducing experimental workload, and enhancing screening efforts for molecules with genuine mechanistic potential for sonodynamic therapy.

Traditional organoids take 3–8 weeks to mature, which makes them impractical for high-throughput discovery and personalized SDT screening. In contrast, our magnetic-field–guided 3D bioprinting platform creates uniform, hypoxic, physiologically relevant microtumors within days, aligning with NIH’s call for organoid-based, non-animal research. These spheroids maintain expression of tumor suppressors (*TP53, PTEN*), oncogenes (*KRAS, MTOR*), and epigenetic regulators (*HDAC3, DNMT3B*), preserving key transcriptional and microenvironmental features while offering unprecedented scalability and speed.

Using these spheroids, we found that ultrasound greatly increased NEO100-driven cell death in glioblastoma, medulloblastoma, meningioma, and breast-to-brain metastasis. In all four tumor types, combining NEO100 with FUS led to dose-dependent decreases in cell viability that were significantly greater than with the drug alone. This broad responsiveness demonstrates NEO100’s natural sonosensitizing ability and shows that ultrasound can be used as a noninvasive, precisely targeted method to enhance drug efficacy. The only exception, LuBM5 lung-to-brain metastasis, showed minimal response to NEO100 with or without ultrasound, indicating that membrane composition, redox buffering capacity, or metabolic state may influence sonodynamic resistance. These findings emphasize the importance of tumor origin and suggest potential directions for future stratification or combination therapies.

Overall, this NAM accelerates the discovery-to-validation process for sonodynamic therapies by incorporating AI-based predictions, rapid 3D tumor fabrication, and ultrasound techniques that can be applied in clinical settings. NEO100 has demonstrated itself as a powerful and broadly effective sonosensitizer tested across various brain tumor types, supporting its potential for future in vivo studies and clinical use. More broadly, this platform signifies a major shift in preclinical testing—one that aligns with NIH’s strategic focus on organoid technologies and reducing reliance on animals, while also enabling fast, scalable, and physiologically relevant research for treatment-resistant brain cancers.

